# Interactions Between the Histone Variant H2Av and the Suppressor of Hairy wing Insulator Influence the DNA Damage Response and Lymph Gland Development in *Drosophila*

**DOI:** 10.1101/2023.05.31.543180

**Authors:** James Ryan Simmons, Justin D. J. Kemp, Mariano Labrador

## Abstract

Genome architecture is regulated by chromatin insulator proteins, and misregulation of insulator function is associated with genome instability and transcriptional regulatory defects in both vertebrates and *Drosophila*. Indeed, mutations of the sole insulator protein in humans, CTCF, are carcinogenic and mutations in the *Drosophila* insulator protein *Suppressor of Hairy wing* [Su(Hw)] lead to chromosomal rearrangements. However, the mechanism that links the DNA damage response and the regulation of transcription with insulator function is not yet understood. Here we show that enrichment of Su(Hw) insulator proteins at insulator sites increases after DNA damage. Additionally, Su(Hw) is necessary for phosphorylation of the histone variant H2Av in response to both UV treatment and X-ray irradiation. The requirement of Su(Hw) for H2Av phosphorylation appears to be tissue-specific, since H2Av is phosphorylated in response to DNA damage also in neurons, where Su(Hw) is not normally expressed. Similarly, we provide evidence that Su(Hw) and H2Av work together to ensure proper development of the lymph gland in *Drosophila* larvae. We show that H2Av regulates formation of the larval lymph gland, and mutation of H2Av causes formation of large melanotic tumors that are rescued by mutation of *su(Hw)* in the *His2Av^810^* mutant background. Double mutants of *su(Hw)^-^* and *His2Av^810^* also form supernumerary lymph gland lobes that are larger than in wild type. Our findings support a role for the Su(Hw) in the DNA damage response through the regulation of H2Av phosphorylation and suggest that Su(Hw) and H2Av also work together to ensure proper development of the lymph gland.

## Introduction

The activity of a diverse array of DNA-binding proteins is responsible for essential chromosomal functions, including transcriptional regulation, progression through the cell cycle, and the DNA damage response. Histone proteins, which bind DNA directly, are key regulatory proteins involved in these processes. Their regulatory nature is largely due to the myriad of post-translational modifications that may be found on many different residues in each histone. These modifications can influence chromatin state, transcriptional output, and the response to DNA damage (Jambhekar et al., 2019). Histone variants provide another mechanism through which cells can regulate chromosomal activity, as seen in the examples of CENP-A, which is enriched at centromeric regions of the genome in many animals, and the mammalian H2A.X, which is phosphorylated in response to DNA double strand breaks and signals to the cell to direct repair machinery to damaged sites (Talbert and Henikoff, 2021).

Another class of DNA-binding proteins are those found in chromatin insulator complexes. Chromatin insulator proteins bind specific sequences found throughout the genome and work in pairs or larger groups to regulate the spread of heterochromatin and to facilitate interactions between gene enhancers and promoters (Chen and Lei, 2019; Dehingia et al., 2022; Phillips-Cremins and Corces, 2013; Schoborg and Labrador, 2014). In vertebrates, transcriptional regulation by insulators works through a mechanism of loop extrusion, in which structural maintenance of chromosome (SMC) proteins cohesin and condensin drive the movement of DNA through the ring complex until encountering an occupied insulator site, setting the boundaries of the loops (Fudenberg et al., 2016); however, this phenomenon has not been observed in *Drosophila* (Matthews and White, 2019). Chromatin insulators also help mitigate the response to DNA damage in human cells (Hilmi et al., 2017) and previous work by our lab characterized the relationship between the phosphorylated form of the H2A variant in *Drosophila*, γH2Av, and insulator proteins. A close association was observed between Su(Hw) insulator proteins and γH2Av throughout the genome by a combination of immunostaining and ChIP-seq analysis (Simmons et al., 2022). Mutation of any component of the Su(Hw) insulator complex was enough to disrupt the interaction of this complex with γH2Av, and *su(Hw)* mutants displayed significantly less γH2Av in polytene chromatin than wild type. The association between insulator proteins and γH2Av was strengthened by treatment with an inhibitor of protein phosphatase 2A (PP2A). Further experiments revealed that γH2Av is a component of insulator bodies that form during osmotic stress and that recovery of the bodies after stress can be prevented by inhibition of PP2A (Simmons et al., 2022). Mutation of *His2Av* suppressed the *ct^6^* phenotype in a sensitized *su(Hw)* heterozygous background, demonstrating that H2Av and Su(Hw) function in the same developmental pathway in the wing. Several questions remained, however, including which kinases might promote this interaction, how lack of H2Av might affect insulator protein recruitment to chromatin, and what the developmental implications are for the interaction between the *His2Av* and *su(Hw)* genes in wild type and mutant larvae. An important question that remains unanswered is whether the interaction between Su(Hw) insulator proteins and H2Av are required for the DNA damage response, for transcriptional regulation under normal conditions, or both.

Here, we first explored the role that Su(Hw) insulator proteins play in the DNA damage response and found that Su(Hw) and Mod(mdg4)67.2 binding to polytene chromosomes increases in response to UV irradiation. Furthermore, H2Av phosphorylation increases on polytene chromosomes in response to UV and X-ray irradiation in a Su(Hw)-dependent manner. Mutation of two key DNA damage response kinases, ATR/*mei-41* and ATM/*tefu*, increases Su(Hw) binding to polytene chromatin under native conditions but prevents the increase of Su(Hw) chromatin binding in response to UV irradiation.

The interaction between Su(Hw) and H2Av is not surprising given the genome instability associated with insulator malfunction in both humans and fruit flies (Hsu et al., 2020; Kemp et al., 2014; Lang et al., 2017). However, the impact that the Su(Hw)-H2Av interaction has on transcriptional regulation under normal conditions remains unclear. This question is important as insulator complexes and H2Av have been independently shown to influence the expression of genes during development (Gambetta and Furlong, 2018; Grigorian et al., 2017; Johnson et al., 2018; Kotova et al., 2011; Ozdemir and Gambetta, 2019; Swaminathan et al., 2005).

To answer this question, we turned to the *Drosophila* larvae to investigate how these proteins work in concert to direct tissue development. We focused on the larval lymph gland, which generates the cells of the circulatory system of *Drosophila*. In larvae, this system is composed of differentiated circulating hemocytes, found in the hemolymph, plus pockets of sessile hemocytes scattered throughout the animal, and the lymph gland which generates most cells of the hemolymph (Banerjee et al., 2019). The lymph gland of a third instar larvae is comprised of three sets of lobes that run from anterior to posterior along the dorsal vessel, which acts as the heart by moving hemocytes through the larvae. The two posterior lobes, the secondary and tertiary lobes, are small and generate few hemocytes at this stage. The anterior-most primary lobe, however, is the largest and contains many undifferentiated prohemocytes in addition to differentiated hematocytes, crystal cells, and lamellocytes (Banerjee et al., 2019). Differentiation of prohemocytes is regulated by Hedgehog signaling generated from cells in the posterior signaling center (PSC). PSC cell identity is determined by expression of *Antennapedia*, the lack of which results in unregulated division of prohemocytes in the primary lobe (Banerjee et al., 2019).

Misregulation of H2Av was recently shown to have a striking phenotype in developing larvae. Individuals homozygous for the *His2Av^810^*null allele, or trans-heterozygous for the *His2Av^810^* null allele paired with a chromosomal deficiency lacking the *His2Av* gene (Df(3R)BSC524), develop large melanotic masses at their posterior ends (Grigorian et al., 2017). The formation of these tumors is correlated with the loss of crystal cells, differentiated plasmatocytes, and the undifferentiated prohemocytes from the primary lobes of the lymph gland. It was proposed that H2Av is necessary for prohemocytes to be competent to respond to Hedgehog signaling from PSC cells, suggesting an inhibited transcriptional response in prohemocytes (Grigorian et al., 2017). Based on our previous results showing extensive colocalization of H2Av with Su(Hw) insulator proteins throughout the genome, we questioned whether interactions between H2Av and Su(Hw) are necessary for proper hematopoiesis in *Drosophila* larvae. Our results reveal a significant role for Su(Hw) in larval hematopoiesis and a genetic interaction with H2Av that influences growth and cell differentiation in the lymph gland. This, together with our findings on the connection between Su(Hw) and H2Av phosphorylation in response to DNA damage, indicates a complex relationship between these two chromatin-binding proteins that is crucial for transcription regulation, development, and genome integrity.

## Materials and Methods

### Fly stocks and husbandry

All stocks were maintained on a standard cornmeal agar fly food medium supplemented with yeast at 20°C; crosses were carried out at 25°C. The following stocks are maintained in our lab and were originally obtained from Dr. Victor Corces (Emory University): *w^1118^*; *su(Hw)^V^*/*TM6B*, *Tb^1^*. The stock *w^1118^*; PBac(RB)*su(Hw)^e04061^*/*TM6B*, *Tb^1^* was obtained from the Bloomington *Drosophila* stock center (BDSC: 18224). The following stocks were provided by our lab: OR, *y^2^w^1^ct^6^*; PBac(RB)*su(Hw)^e04061^*/*TM6B*, *Tb^1^* (derived from BDSC: 18224), *y^2^w^1^ct^6^*; P{Su(Hw)::eGFP}/Cyo; PBac(RB)*su(Hw)^e04061^*/*TM6B*, *Tb^1^*.

### Antibodies

Rabbit polyclonal IgG antibodies against Su(Hw), Mod(mdg4)67.2, and CP190 were previously generated by our lab (Schoborg et al., 2013; Wallace et al., 2010). An antibody against the phosphorylated form of H2Av (UNC93-5.2.1) (Lake et al., 2013) was obtained from the Developmental Studies Hybridoma Bank, created by the NICHD of the NIH and maintained at The University of Iowa, Department of Biology, Iowa City, IA 52242. These antibodies were all diluted 1:1 in glycerol (Fisher Scientific, BP229-1, lot 020133) and used at a final dilution of 1:200. Secondary antibodies were all diluted 1:1 in glycerol and used at a final dilution of 1:200. The following secondary antibodies were used in this study: Alexa Fluor 594 goat anti-rabbit (Invitrogen, A-111037, lot 2079421), Alexa Fluor 488 donkey anti-rabbit (Invitrogen, A-21206, lot 1834802), Alexa Fluor 488 goat anti-mouse (Invitrogen, A-11001, lot 1858182), and a Texas Red anti-mouse (Jackson Laboratories).

### Immunostaining of larval tissues

Wandering third instar larvae were dissected in PBS. Tissues were immediately placed into fixative (4%, para-formaldehyde (Alfa Aesar, 43368, lot N13E011), 50% glacial acetic acid (Fisher Scientific, A38-212, lot 172788) on a coverslip for one minute. Samples were squashed by lowering a slide on top of the sample then turning it over, placing it between sheets of blotting paper, and hitting the coverslip firmly with a small rubber mallet. Slides were dipped in liquid nitrogen, coverslips were removed, and samples were incubated in blocking solution (3% powdered nonfat milk in PBS + 0.1% IGEPAL CA-630 (Sigma-Aldrich, 18896, lot 1043) for 10 minutes at room temperature. The slides were dried and incubated with primary antibodies overnight at 4°C in a box humidified with wet paper towels. The next day, slides were washed twice in PBS + 0.1% IGEPAL CA-630 before incubation with secondary antibodies for three hours in the dark at room temperature. Slides were washed twice in PBS + 0.1% IGEPAL CA-630, treated with DAPI solution of 0.5 μg/mL (ThermoFisher, D1306) for one minute, and washed one more time in PBS alone.

Samples were mounted with Vectashield antifade mounting medium (Vector Laboratories, H-1000, lot ZF0409) and coverslips were sealed with clear nail polish. All microscopy for immunostaining was performed on a wide-field epifluorescent microscope (DM6000 B; Leica Microsystems) equipped with a 100X (NA 1.35) oil immersion objective and a charge-coupled device camera (ORCA-ER; Hamamatsu Photonics). Image acquisition was performed using SimplePCI (v6.6; Hamamatsu Photonics). Image manipulation was performed in FIJI (Schindelin et al., 2012); all contrast adjustments are linear. Images were further processed in Adobe Photoshop CS5 Extended, Version 12.0 x64. Figures were assembled in Adobe Illustrator CS5, Version 15.0.0. Statistical analyses were performed in GraphPad Prism version 8.0.0 (224) (GraphPad Software, San Diego, CA).

### Fluorescence intensity and colocalization analysis

Images were analyzed for the intensity of each channel (as a proxy for the amount of each protein)) using a macro script in FIJI (Schindelin et al., 2012). The DAPI channel was used to automatically generate non-biased ROIs for each cell, which were then manually curated for extra precision. A rolling-ball background subtraction algorithm was used for all images. Intensity measurements were made using the measure function. Numerous chromosomal images were collected from each sample. All acquisition parameters were kept constant between slides within each experiment.

### Western blotting

Tissues were dissected from wandering third instar larvae in 1X PBS. Tissues were collected in RIPA buffer (150 mM NaCl, 1% NP-40, 0.5% sodium deoxycholate, 0.1% SDS, 50 mM Tris-HCl, pH 8.0) supplemented with protease inhibitor (A32953, Thermo Scientific) and a phosphatase inhibitor (A32957, Thermo Scientific). Samples were homogenized in a 500 μL PCR tube with a motorized pestle, mixed with 5X loading buffer (0.5 M Tris-HCl, pH6.8, 5% SDS, 0.0125% bromophenol blue, 12.5% β-mercaptoethanol, 50% glycerol) and 15% per volume β-mercaptoethanol, and boiled for 10 minutes. Protein samples were separated on 8% polyacrylamide gels and transferred onto 0.45 μm PVDF membranes. Membranes were blocked in a solution of 20% Odyssey Blocking Buffer (LI-COR, 927-50000) in TBS at room temperature for one hour. Membranes were incubated in primary antibodies diluted in TBST at 4 °C overnight. Primary antibodies used in these western blots include the mouse α-γH2Av (1:5,000) and our rabbit polyclonal α-H2Av antibody (1:500). Next, membranes were washed four times in TBST, 5 minutes each wash. Membranes were then incubated in secondary antibodies diluted in TBST at room temperature for one hour. Secondary antibodies used in these western blots include goat α-rabbit 680 RD (1:10,000) and donkey α-mouse 800 CW (1:10,000). Membranes were washed four times in TBST followed by two washes in TBS, 5 minutes each wash. Blots were imaged on an Odyssey CLx scanner (LI-COR). Image files were analyzed using FIJI (Schindelin et al., 2012).

### UV and X-ray irradiation

For UV-irradiation, third instar larvae were collected in a small volume of 1X PBS on a glass dissecting dish. Larvae were exposed to varying doses of UV irradiation in a UV Stratalinker 2400 (Stratagene), then allowed to recover for three hours at room temperature in a vial of fly food before processing each sample. For X-ray irradiation, third instar larvae were collected in a plastic chamber and irradiated at different doses in an RS2000 Biological Irradiator (Rad Source Technologies, inc.) at a rate of 2 Gy/min. Irradiated larvae were placed in a vial of fly food and larval samples were processed after three hours of recovery. For western blots, tissues from 10 individual larvae were collected and pooled as one sample. For the lethality assays, 10 larvae of each genotype were irradiated. This was performed in triplicate for each dose.

### Generation of *su(Hw)^-^ His2Av^810^* double mutant genotypes

To generate double mutants, flies with null mutations of *su(Hw)^e04061^*and *His2Av^810^* in the third chromosome were crossed (See appendix, Figure A1). Female F1 progeny in which recombination between the two mutant chromosomes happened were crossed against the *His2Av^810^*/*TM6B* mutant background and potential recombinant chromosomes were isolated in F2 males over the *TM6B* balancer. F2 males were selected for the presence of the *su(Hw)^e04061^* mutation based on eye color and used to establish isogenic lines after two more generations of crossing. As a control for potential second-site mutations on the *su(Hw)^e04061^* chromosome, double mutants were also generated using the *su(Hw)^V^* mutations. Flies with null mutations of *su(Hw)^V^* and *His2Av^810^* were crossed in the background of the *y^2^ct^6^ gypsy* insulator mutations (See appendix, Figure A2). Female F1 progeny in which recombination between the two mutant chromosomes happened were crossed against the *su(Hw)^e04061^*/*TM6B* mutant background and potential recombinant chromosomes were isolated in F2 males over the *TM6B* balancer. F2 males were selected for the presence of the *su(Hw)^V^* mutation based on the rescue of the *y^2^ct^6^* mutant phenotype and used to establish isogenic lines after two more generations of crossing. All mutant lines were genotyped by PCR to determine the presence or absence of the *His2Av^810^* mutation.

### Crystal cell detection and lymph gland phenotyping

Crystal cells in wandering third instar larvae were visualized by incubating between 5-10 larvae in 40 μL 1X PBS buffer in a 200 μL PCR tube in a thermocycler set to 70 °C for 10 minutes. Samples were left overnight at 4 °C to increase the intensity of the melanization product for easier detection of crystal cells and lymph glands. Larvae were placed in halocarbon oil 700 (Sigma, H8898) and photographed using a stereomicroscope (MZ16 FA; Leica Microsystems) equipped with a CCD color camera (DFC420; Leica Microsystems). A 150 Watt white light source (KL 1500 LCD; Leica Microsystems) set to a color temperature of 3,000 K was used for illumination. Images were collected using Leica Application Suite (Version 2.4.0 R1; Leica Microsystems). Image analysis was performed in FIJI (Schindelin et al., 2012) by counting the number of crystal cells visible on the dorsal side of each larvae. The frequency at which lymph glands become visible after this treatment was recorded, as were measurements of the number of lobes in each lymph gland and the size of each lobe.

### Sterile wound assay

To determine whether mutant larvae had functional innate immune responses, a sterile wound assay was performed. Wandering third instar larvae were selected, washed briefly in 1X PBS buffer and 70% ethanol, and placed on a soft pad constructed of black construction paper (for contrast) taped to a folded-up paper towel. Larvae were stabbed on the dorsal side along the midline approximately between abdominal segments A3 to A4 (Galko and Krasnow, 2004) using an insulin syringe needle (Beckton, Dickinson, and Company, Micro-fine IV 28G1/2, 0.36 mm diameter). In some cases, the needle penetrated all the way through the ventral side of the larvae and clots were visible on both sides. Larvae were incubated in a vial of fly food overnight at 25 °C before scoring. Larvae that died immediately after stabbing due to unintentional damage were discarded from the count.

### Insulator protein enrichment at the *Antp* gene

Publicly available ChIP-Seq datasets for Su(Hw), dCTCF and CP190 were obtained from (Wood et al., 2011) and for H2Av from (Li et al., 2016) were used to map binding sites of the insulator proteins Su(Hw), CP190, dCTCF, and γH2Av. The sequencing data was uploaded to the Galaxy web platform, and the public server at usegalaxy.org was used for analysis (Afgan et al., 2018). Briefly, the FastQ datasets from NCBI were mapped with Bowtie2 to produce BAM files (Langmead and Salzberg, 2012) and Bigwig files were generated by applying Bamcoverage to BAM files. Duplicate and unmapped reads were filtered out with SAM tools. The IGV genome browser (igv.org/app/) (Robinson et al., 2011), was utilized to visualize peak profiles and generate peak profile figures using *Drosophila* dm6 as the reference genome.

## Results

### Polytene chromosomes require Su(Hw) to respond to DNA damage

In order to determine whether the interaction between chromatin insulator proteins and H2Av is related to the histone variant’s role in the DNA damage response, third instar larvae were treated with UV irradiation and allowed three hours recovery. Immunostaining of salivary gland polytene chromosomes revealed a significant increase in the amount of H2Av phosphorylation in the wild type OR strain after UV treatment when compared to the unexposed control (Figure 1 C, D), indicating the presence of DNA damage. Additionally, the immunostaining intensity of Su(Hw) protein on polytene chromatin increased significantly after UV irradiation of wild type larvae (Figure 1 C, D). However, this finding was not replicated in diploid cells in the larval brain that express little *su(Hw)*. Even though increased H2Av phosphorylation was observed after UV treatment (Figure 1 A, B), no significant increase in Su(Hw) signal was seen, suggesting this response is tissue specific and limited to cells that express Su(Hw). Despite this, a sensitivity to UV irradiation was displayed by *su(Hw)^e04061^* homozygous mutants in a lethality test performed at various doses. *su(Hw)^e04061^* mutants experience significant developmental defects and associated lethality under control conditions (Figure 1E). Larvae homozygous for this mutation show 100% lethality at a UV irradiation dose of 25 mJ/cm^2^, whereas *su(Hw)^e04061^* heterozygous mutants and the OR wild type control are still viable at this dose, albeit at lower rates (Figure 1E). This suggests that mutation of *su(Hw)* impairs the larvae’s ability to respond to DNA damage caused by UV irradiation.

**Figure 1.**
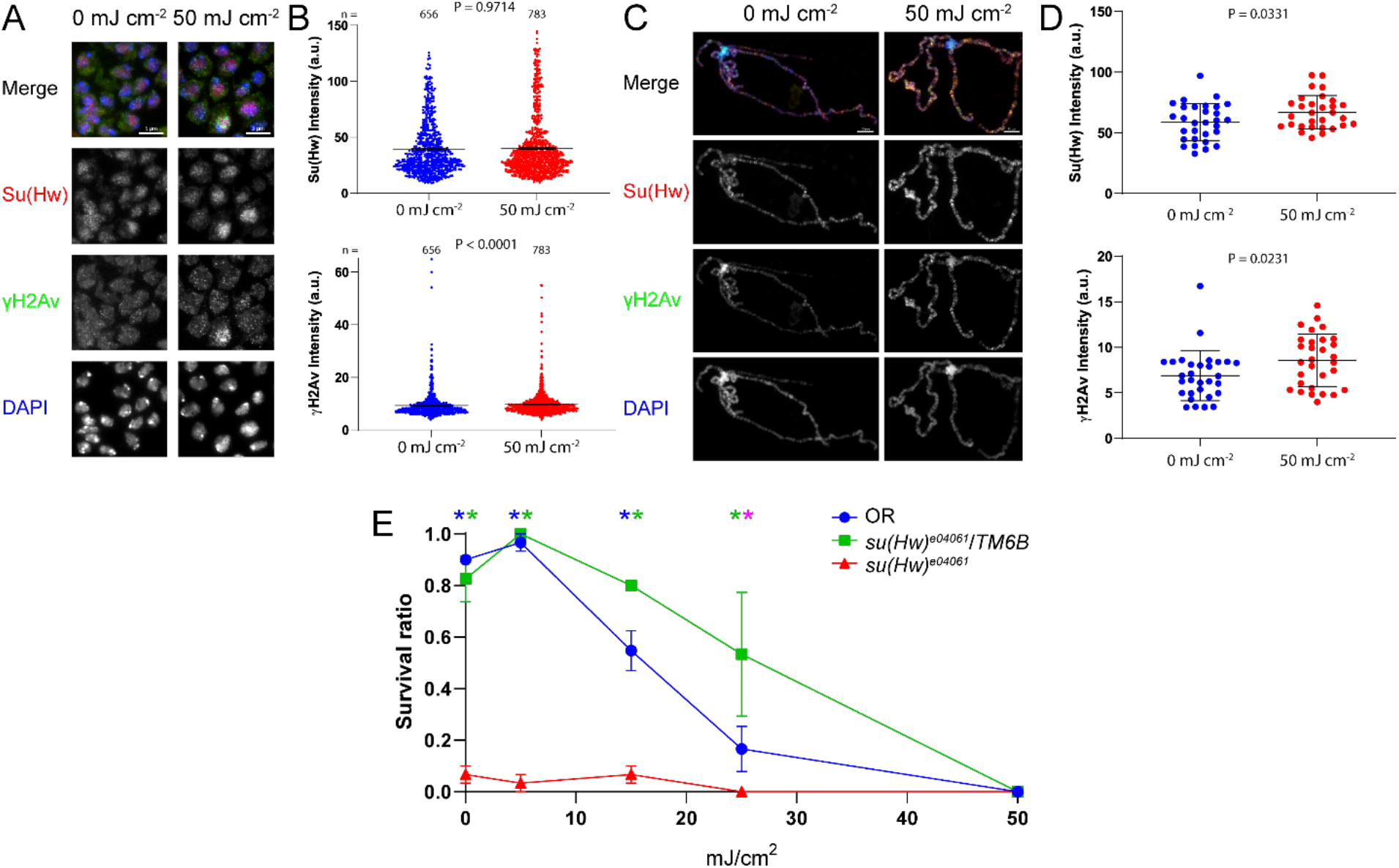
Su(Hw) and H2Av phosphorylation increase in polytene chromatin in response to UV irradiation. **A.** Immunostaining results for Su(Hw) and γH2Av antibodies in neural cells with and without UV treatment. The scale bars are 5 μm each. **B.** Fluorescent intensities of the Su(Hw) and γH2Av antibody signals in neural cells. P-values were determined using unpaired ANOVA with Dunn’s multiple comparison test. **C.** Immunostaining results for Su(Hw) and γH2Av antibodies in polytene chromatin with and without UV treatment. The scale bars are 10 μm each. **D.** Fluorescent intensities of the Su(Hw) and γH2Av antibody signals in polytene chromatin. Each dot in B and D represents an individual cellular genome. P-values were determined using unpaired Welch’s T-tests. **E**. Lethality assay for UV irradiation. Blue asterisks represent a significant difference between OR and *su(Hw)^e04061^*, green asterisks represent a significant difference between *su(Hw)^e04061^*/*TM6B* and *su(Hw)^e04061^*, and magenta asterisks represent a significant difference between OR and *su(Hw)^e04061^*/*TM6B*. P-values were determined using a two-way ANOVA with Tukey’s multiple comparisons test at each dosage, with values below 0.05 considered significant. Error bars represent one standard error of the mean.

The level of Su(Hw) protein on polytene chromatin was significantly decreased in the *su(Hw)^e04061^* mutant larvae compared to the wild type, as was the level of Mod(mdg4)67.2, which depends on Su(Hw) for recruitment to chromatin (Figure 2 A,B). Like Su(Hw), Mod(mdg4)67.2 levels increase significantly in polytene chromosomes in response to UV irradiation (Figure 2 A,B). This increase in Mod(mdg4)67.2 is dependent on Su(Hw), as no change in Mod(mdg4)67.2 levels is seen in the *su(Hw)^e04061^*mutant after UV treatment (Figure 2 A,B). More significantly, the increase in H2Av phosphorylation that occurs in wild type polytene chromosomes in response to UV irradiation is abolished in the *su(Hw)^e04061^* mutant (Figure 2 A,B), indicating that H2Av phosphorylation upon DNA damage depends on a mechanism mediated by Su(Hw).

**Figure 2.**
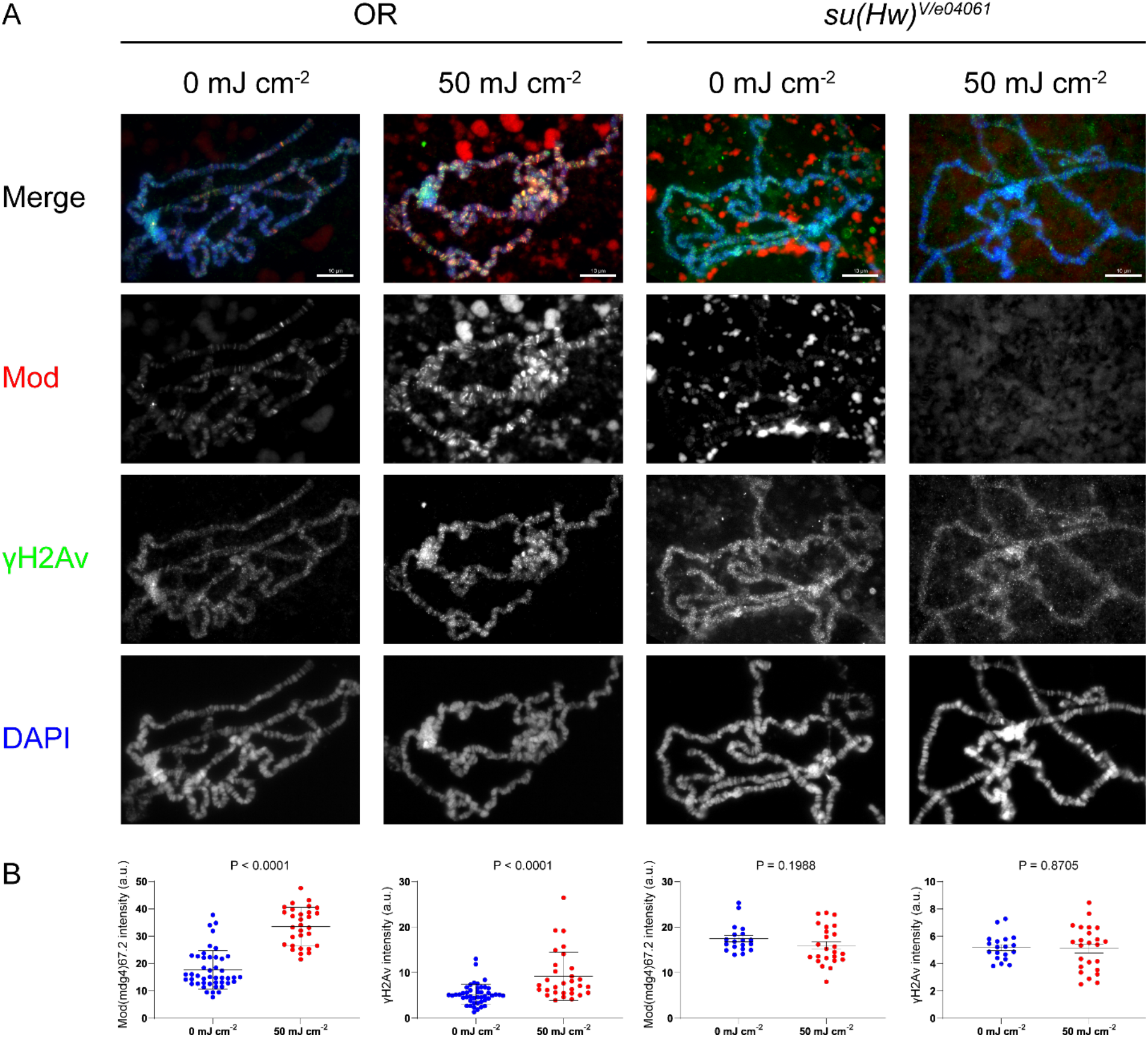
Su(Hw) is necessary for H2Av phosphorylation in polytene chromatin after UV irradiation. **A.** Immunostaining results for Mod(mdg4)67.2 (abbreviated as Mod) and γH2Av antibodies on polytene chromosomes with and without UV treatment. The scale bars are 10 μm each. **B.** Fluorescent intensities of the Mod(mdg4)67.2 and γH2Av antibody signals. Each dot represents a single polytene genome. P-values were determined using unpaired Welch’s T-tests.

To determine if Su(Hw) is involved in the cellular response to ionizing radiation, third instar larvae were subjected to a 10 Gy dose of X-rays and allowed to recover for four hours. Immunostaining of diploid cells from the larval brain of unexposed control samples revealed a significantly lower degree of H2Av phosphorylation in the *su(Hw)^e04061^*mutant as compared to wild type (Figure 3 A, B), a finding consistent with previous results in polytene chromatin (Simmons et al., 2022). However, upon X-ray irradiation, phosphorylated H2Av immunostaining levels increased significantly in both the wild type larvae and the *su(Hw)^e04061^* mutant (Figure 3 A, B). Likewise, western blotting of larval brains demonstrates a similar degree of H2Av phosphorylation upon X-ray treatment in the wild type and *su(Hw)^e04061^* mutant larvae (Figure 3 C). Immunostaining reveals a near complete absence of Su(Hw) in the *su(Hw)^e04061^* mutant, and there is no change in Su(Hw) staining levels in wild type after X-ray irradiation (Figure 3 A, B). These findings suggest that Su(Hw) is not necessary for H2Av phosphorylation in neural cells in response to ionizing radiation.

**Figure 3.**
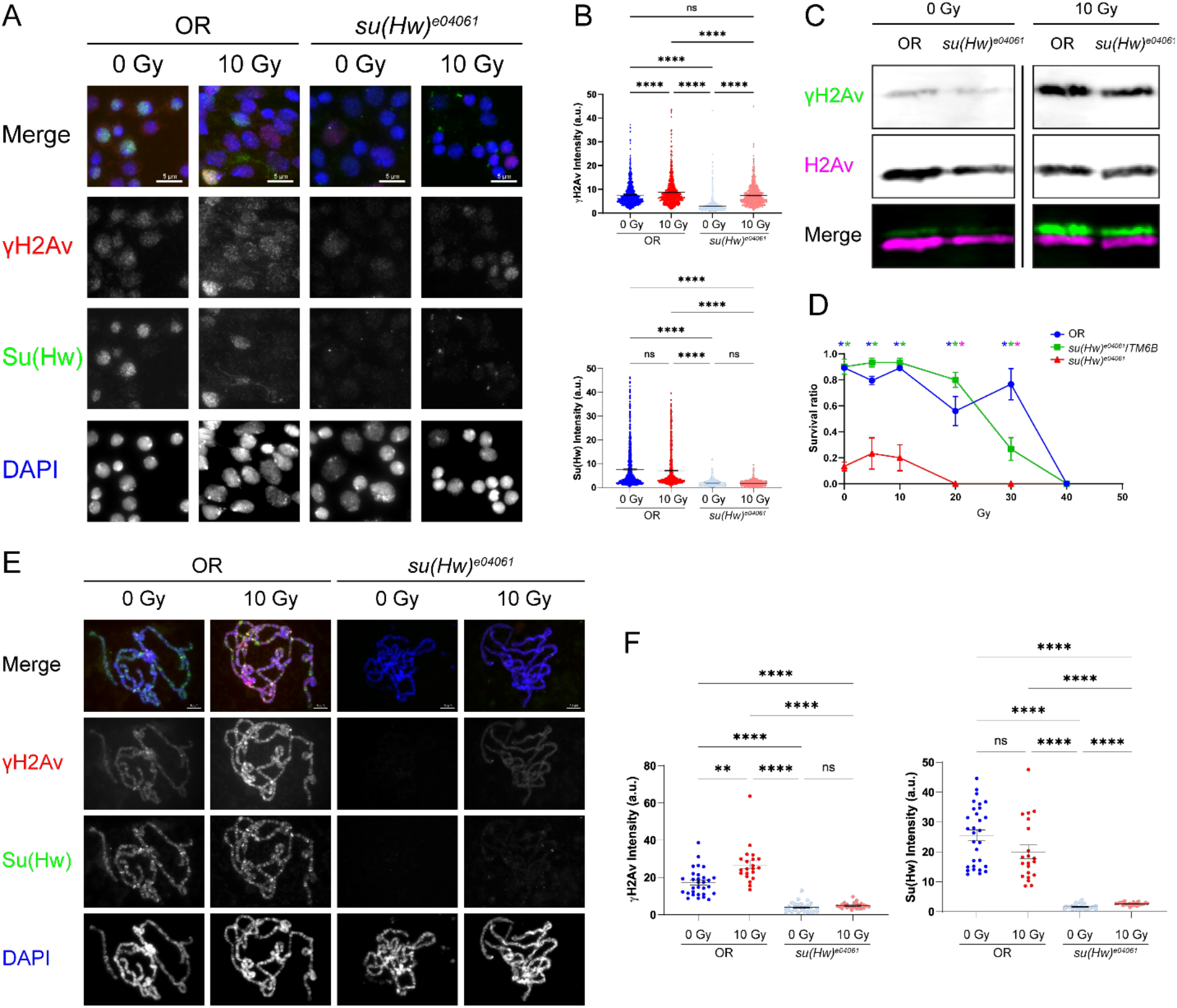
Su(Hw) is necessary for the H2Av phosphorylation in polytene chromatin after X-ray irradiation. **A.** Immunostaining results for Su(Hw) and γH2Av antibodies in neural cells with and without UV treatment. The scale bars are 5 μm each. **B.** Fluorescent intensities of the Su(Hw) and γH2Av antibody signals. P-values were determined using unpaired ANOVA with Dunn’s multiple comparison test. **C.** Western blotting results for central nervous systems from control and X-ray irradiated larvae. **D.** Lethality assay for X-ray irradiation. Blue asterisks represent a significant difference between OR and *su(Hw)^e04061^*, green asterisks represent a significant difference between *su(Hw)^e04061^*/*TM6B* and *su(Hw)^e04061^*, and magenta asterisks represent a significant difference between OR and *su(Hw)^e04061^*/*TM6B*. P-values were determined using a two-way ANOVA with Tukey’s multiple comparisons test at each dosage, with values below 0.05 considered significant. Error bars represent one standard error of the mean. **E**. Immunostaining results for Su(Hw) and γH2Av antibodies in polytene chromatin with and without UV treatment. The scale bars are 5 μm each. **F.** Fluorescent intensities of the Su(Hw) and γH2Av antibody signals. P-values were determined using unpaired ANOVA with Dunnett’s T3 multiple comparison test. Each dot in B and E represents a single polytene genome. ** P < 0.01, **** P < 0.0001.

Immunostaining in polytene chromosomes shows the expected pattern for Su(Hw), with significantly lower levels in *su(Hw)^e04061^* mutants compared to the wild type control (Figure 3 E, F). Like diploid cells, the level of nuclear Su(Hw) does not change in polytene chromosomes after X-ray treatment (Figure 3 E, F). As expected, the level of H2Av phosphorylation increases significantly in polytene chromosomes after X-ray treatment (Figure 3 E, F). Unlike diploid cells, however, this response is inhibited in *su(Hw)^e04061^*mutants, which show little γH2Av after X-ray irradiation (Figure 3 E, F). This is similar to the response seen after UV irradiation in the *su(Hw)^e04061^*mutants, hinting at a mechanism in which Su(Hw) is required for H2Av phosphorylation upon various forms of DNA damage in salivary gland cells. A lethality assay was performed on third-instar larvae using a range of X-ray radiation doses. While many of the *su(Hw)^e04061^*homozygous mutants fail to survive even in the absence of radiation, a dose of 20 Gy is enough to produce full lethality (Figure 3D). In contrast, most of the *su(Hw)^e04061^* heterozygous mutants and the OR wild-type control larvae survived after a dose of 20 Gy, with survival for both genotypes falling to zero around 40 Gy (Figure 3D). These findings further support the notion that Su(Hw) is required for DNA damage repair in response to ionizing radiation.

### Simultaneous mutation of ATR and ATM disrupts Su(Hw) binding dynamics to chromatin

Based on our previous results, phosphorylation of H2Av stabilizes insulator protein complexes in polytene chromatin. Several kinases have been reported to phosphorylate H2Av, with the *Drosophila* homologs for ATR (*mei-41*) and ATM (*tefu*) being the key examples (Joyce et al., 2011). We therefore sought to determine the effect of mutation of each of these kinases by immunostaining polytene chromatin for Su(Hw). We found no difference in the amount of Su(Hw) bound to chromatin in either single mutant (*mei-41^D5^* and *tefu^atm-3^*); however, there was a significant increase in the amount of Su(Hw) in the double mutant (Figure 4 A, B). In addition, we measured the effect of these mutations on H2Av phosphorylation in polytene chromatin. We found a significant increase in H2Av phosphorylation in the ATR/*mei-41^D5^* mutant but not the ATM/*tefu^atm-3^*mutant (Figure 4 A, B). The double mutant also showed significantly more H2Av phosphorylation than wild type, though there was no difference between the double mutant and the ATR/*mei-41* single mutant, suggesting the difference may be due to the *mei-41* mutation alone (Figure 4 A, B).

**Figure 4.**
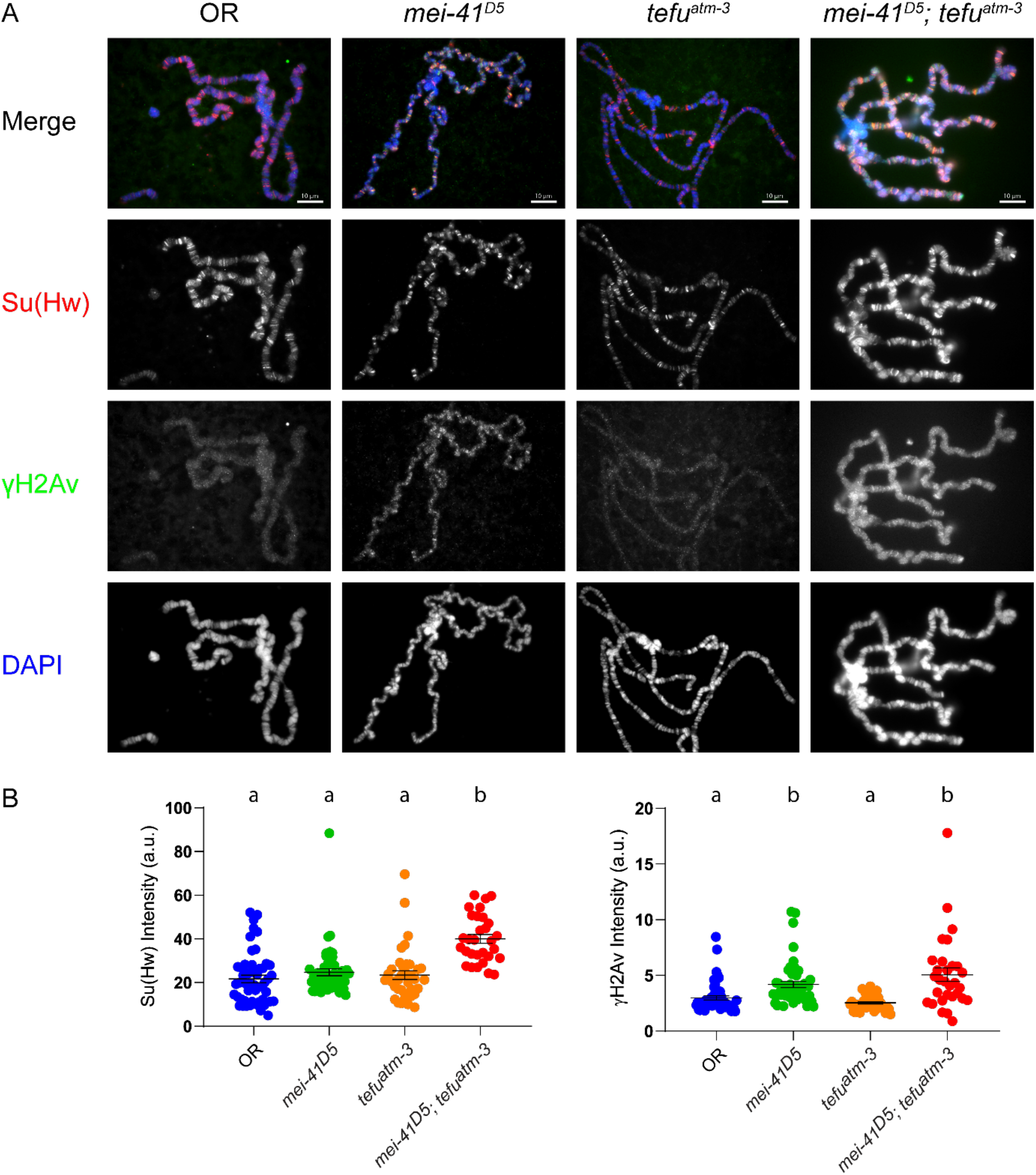
Simultaneous mutation of ATR and ATM increases Su(Hw) bound to chromatin. **A.** Immunostaining results for Su(Hw) and γH2Av antibodies on polytene chromosomes. The scale bars are 10 μm each. **B.** Fluorescent intensities of the Su(Hw) and γH2Av antibody signals. Each dot represents a single polytene genome. Letters above the graphs represent statistical groupings based on a one-way ANOVA with Dunn’s multiple comparisons test with significances at P < 0.05.

Due to the increase in Su(Hw) binding in polytene chromosomes in the double mutant (Figure 4) and after UV irradiation (Figure 1), we questioned what the response to UV irradiation would be in these mutants. Su(Hw) immunostaining intensity increased after UV treatment in the ATR/*mei-41^D5^* mutant but not the ATR/*mei-41^D5^* ATM/*tefu^atm-3^*double mutant (Figure 5 A, B). The lack of response in the double mutant may be due to natively high levels of Su(Hw) observed in the strain (Figure 4 A, B). H2Av phosphorylation increased in response to UV irradiation in both the ATR/*mei-41^D5^*mutant and the ATR/*mei-41^D5^* ATM/*tefu^atm-3^* double mutant (Figure 5 A, B), suggesting at least one other kinase may be acting redundantly with ATR and ATM in the UV response.

**Figure 5.**
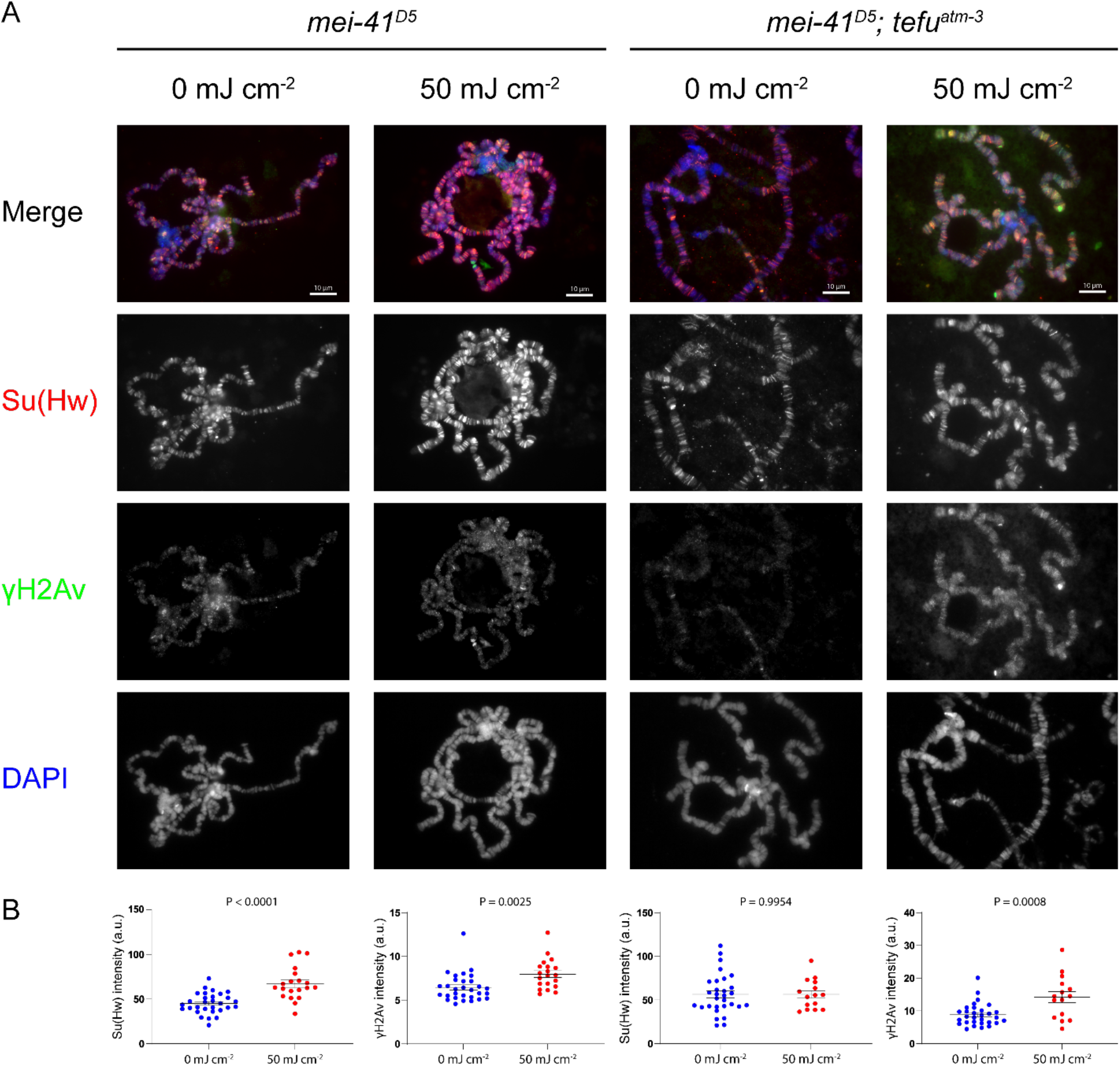
Su(Hw) enrichment in polytene chromatin after UV irradiation depends on ATR and ATM. **A.** Immunostaining results for Su(Hw) and γH2Av antibodies on polytene chromosomes with and without UV treatment. The scale bars are 10 μm each. **B.** Fluorescent intensities of the Su(Hw) and γH2Av antibody signals. Each dot represents a single polytene genome. P-values were determined using unpaired Welch’s T-tests.

### Mutation of *su(Hw)* suppresses the *His2Av^810^* melanotic tumor phenotype

*Drosophila* H2Av is involved in both DNA damage repair and in transcriptional regulation. As we previously explored the relationship between Su(Hw) and H2Av phosphorylation in insulator protein mutants, we next questioned how insulator activity and binding are affected by mutation of *His2Av*. Heterozygous mutation of *His2Av* in a heterozygous *su(Hw)* mutant background partially rescues defects observed in the wing margins of *ct^6^* mutants (Simmons et al., 2022). Based on this, and on previous immunostains showing extensive colocalization between Su(Hw) insulator components and γH2Av (Simmons et al., 2022), we examined binding of insulator proteins to polytene chromatin in the *His2Av^810^* null background. Quantification of immunostaining signals revealed a slight but statistically significant increase in Su(Hw) binding when comparing *His2Av^810^* homozygotes with *His2Av^810^*/*TM6B* heterozygotes (Supplementary Figure 1 A). No difference was observed in the binding of the two other key components of the Su(Hw) insulator complex, Mod(mdg4)67.2 and CP190 (Supplementary Figure 1 B, C), implying the effect of H2Av mutation on insulator binding to polytene chromatin is limited to Su(Hw).

Our next goal was to determine the effect of mutating insulator proteins on the phenotype of *His2Av* mutants. To further explore this relationship between Su(Hw) and H2Av, we generated the double mutant genotype *su(Hw)^-^ His2Av^810^* using established null mutations of *su(Hw)*. As both genes reside on the third chromosome, double mutant generation was performed by taking advantage of meiotic recombination of chromosomes that is largely limited to female *Drosophila* (see methods and the appendix for details). Double mutant *su(Hw)^-^ His2Av^810^* strains were confirmed by PCR and immunostained for insulator proteins and γH2Av. Staining for Su(Hw), Mod(mdg4)67.2, and γH2Av were all significantly reduced in the *su(Hw)^e04061^ His2Av^810^* mutants when compared to heterozygotes (Supplementary Figure 2 A, B, and E); however, CP190 and CTCF staining were unaffected (Supplementary Figure 2 C, D).

*His2Av^810^* null mutants were recently described to form large melanotic masses in their posterior ends as third instar larvae (Grigorian et al., 2017). Inspection of the new double homozygous *su(Hw)^-^ His2Av^810^* genotypes revealed a nearly complete rescue of the melanotic tumor phenotype presented by *His2Av^810^* homozygotes (Figure 6), implying a genetic interaction between the two genes that influences the outcome of the fly hematopoiesis. Whereas the vast majority of *His2Av^810^* homozygous larvae developed large melanotic tumors in their posterior ends, none of the examined *su(Hw)^e04061^ His2Av^810^* homozygous larvae showed such masses. This result was replicated in the other double homozygous mutant *su(Hw)^V/e04061^ His2Av^810^*, suggesting that the rescue is most likely due to mutation of the *su(Hw)* gene specifically and not due to a second-site mutation on the same chromosome (Figure 6). Crossing the double mutants against the single *His2Av^810^* mutant allowed us to monitor how a partial reduction in the dose of *su(Hw)* affects formation of melanotic tumors in the *His2Av^810^* homozygote. Unlike the double homozygote, heterozygote *su(Hw)* mutant larvae generated from this cross showed melanotic masses but at a significantly lower frequency than the *His2Av^810^* mutants alone (Figure 6). These results demonstrate that these masses form in the presence of Su(Hw), and that reducing the amount of Su(Hw) available in the cell is associated with decreased tumor size and frequency in a dose-dependent manner.

**Figure 6.**
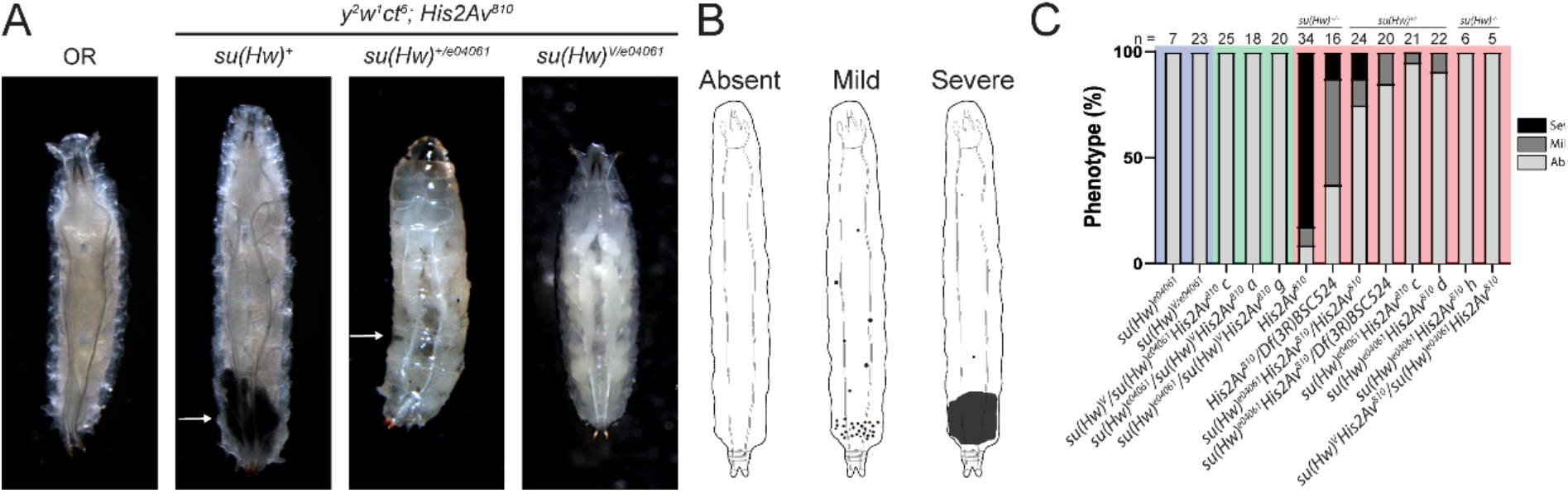
Mutation of *su(Hw)* suppresses the *His2Av^810^* melanotic tumor phenotype. **A.** Representative pictures of select genotypes showing a dorsal view. The presence of black melanotic masses is indicated by white arrows. **B.** Illustration showing presentation of the melanotic tumor phenotype in wandering third-instar larvae (representing each of the phenotypic classes used for tumor scoring: absent, mild, and severe). **C.** Quantification of tumor phenotype scoring across each genotype examined. Within the bars the light grey color indicates larvae lacking tumors, the dark grey indicates a mild phenotype, and the black represents a severe phenotype. The blue background in the graph indicates homozygosity for wild type *His2Av*, while green indicates heterozygosity for *His2Av^810^*, and red indicates homozygosity for *His2Av^810^*. Sample size is shown above each genotype.

### Double mutant *su(Hw)* and *His2Av^810^* larvae exhibit overgrowth of lymph glands

To further characterize the relationship between Su(Hw) and H2Av in the hematopoietic lineage, third instar larvae were subjected to heat treatment at 70°C. This heat treatment triggers catalysis of the prophenoloxidase enzymes found in crystal cells, resulting in cytoplasmic melanin production and subsequent darkening of the cell (Banerjee et al., 2019). This provides a convenient means to visualize the numbers and distribution of crystal cells in heat-treated larvae (Figure 7 A). Heat-treated wild type larvae show around 50 clusters of crystal cells visible from the dorsal side, while *su(Hw)^e04061^* homozygous larvae display significantly more crystal cell clusters with around 200 visible per larvae (Figure 7 B, C). *His2Av^810^* homozygous larvae show similar numbers of crystal cell clusters as wild type; however, this mutant was reported to have reduced lymph glands and exhibits large melanotic tumors that contain many crystal cells (Grigorian et al., 2017), obfuscating the interpretation of this genotype. *su(Hw)^e04061^ His2Av^810^* homozygous double mutant larvae generate significantly more clusters of crystal cells as the wild type but a similar number compared to the *su(Hw)^e04061^* single mutant (Figure 7 B, C).

**Figure 7.**
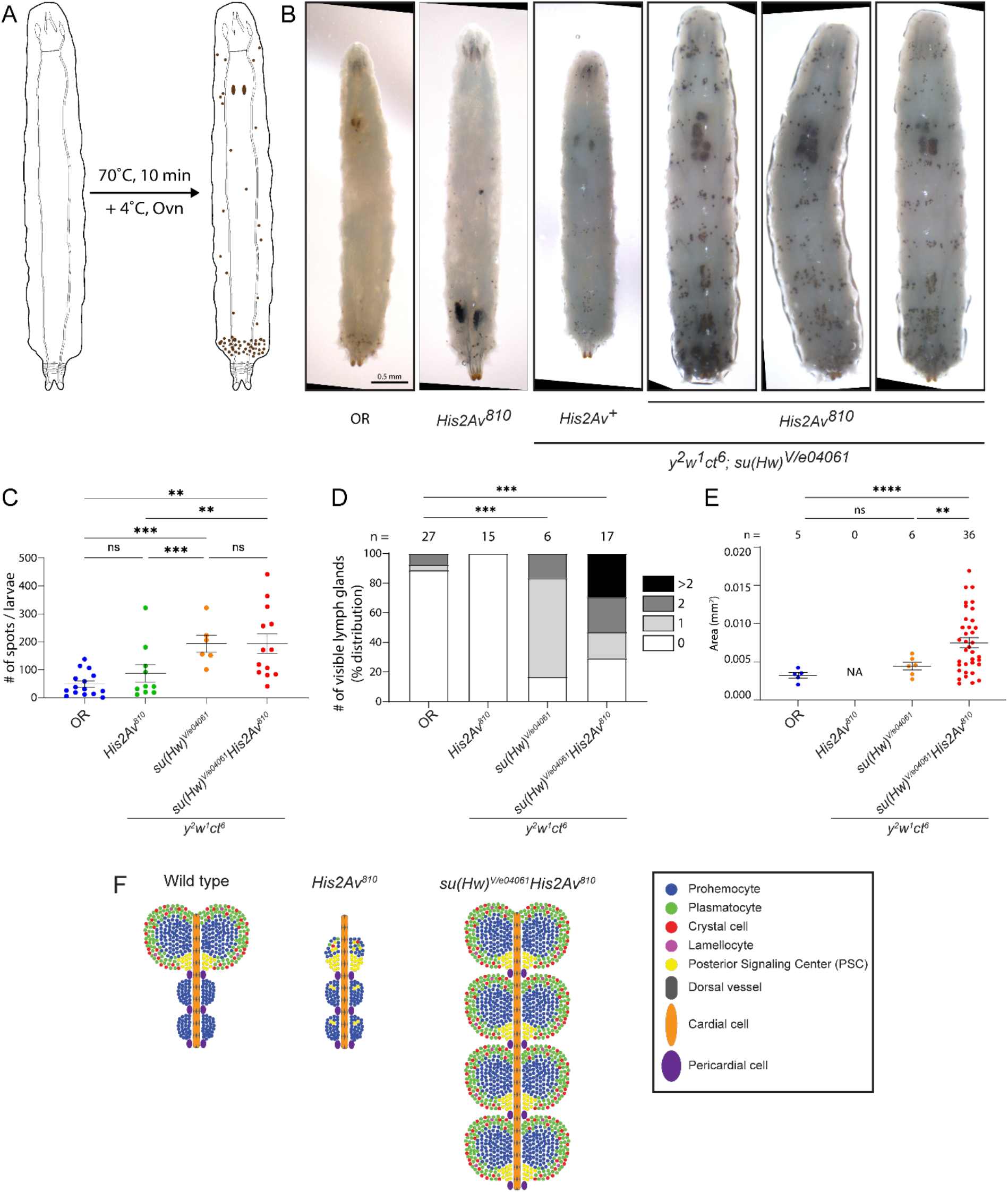
*su(Hw)^-^His2Av^810^* larvae exhibit overgrowth of lymph glands. **A.** Illustration showing the heat treatment assay, which involves incubating larvae at 70 °C and monitoring for melanization of crystal cells and lymph glands. **B.** Representative examples of melanized cells and lymph glands. The *y^2^w^1^ct^6^*; *su(Hw)^V/e04061^His2Av^810^* mutant genotype shows larger lymph glands with supernumerary lobes. The scale bar represents 0.5 mm. **C.** Quantification of melanized spots observed after heat treatment for each genotype. P-values were determined using an ordinary one-way ANOVA with Tukey’s multiple comparisons test. **D.** Number of lymph gland nodes per larvae observed in each genotype. P-values were determined using pairwise Chi-squared tests. **E.** Measurements of lymph gland lobe size in different genotypes. P-values were determined using ANOVA with Dunnett’s multiple comparisons. * P < 0.05, ** P < 0.01, *** P < 0.001, **** P < 0.0001. **F.** Model of lymph gland development in wild type, *His2Av^810^*, and *su(Hw)^V/e04061^His2Av^810^* larvae.

In addition to the circulating crystal cells that become visible upon heat treatment, the lymph glands themselves sometimes become visible due to the presence of crystal cells (Figure 7 A). The frequency of observable lymph glands after heat treatment is significantly higher in the *su(Hw)^e04061^ His2Av^810^* mutants than the wild type (Figure 7 D). Shockingly, the lymph glands in *su(Hw)^e04061^ His2Av^810^* double mutants were significantly overdeveloped compared to those in wild type larvae, often showing supernumerary lobes (Figure 7 B, D). Indeed, some larvae in the *su(Hw)^e04061^ His2Av^810^* homozygous mutant displayed up to eight large lymph gland lobes, and the average size of lymph gland lobes was significantly larger in the *su(Hw)^e04061^ His2Av^810^* double mutant than wild type (Figure 7 B, D-F). Supernumerary lymph gland lobes were also observed at a low frequency in larvae homozygous for *su(Hw)^e04061^* and heterozygous for *His2Av^810^*(Figure 7 D).

Since immune system development in *Drosophila* larvae depends largely upon hematopoiesis in the lymph gland, we tested the innate immune response in these larvae using a sterile wound assay. This involves stabbing larvae using an ultrafine sterile needle and monitoring surviving larvae for clot formation and melanization at the site of wounding (Supplementary Figure 3 A). This assay revealed that, like the wild type strain, *su(Hw)^e04061^ His2Av^810^* double mutants are competent to form a clot in response to sterile injury (Supplementary Figure 3 B, C). *su(Hw)^e04061^*single mutants also form clots in response to sterile wounds; however, *His2Av^810^*mutants form clots at a slightly lower frequency compared to wild type (Supplementary Figure 3 C), likely a result of the hematopoietic phenotypes reported previously (Grigorian et al., 2017). Despite the rescue of the *His2Av^810^* hematopoietic phenotypes observed in the *su(Hw)^e04061^ His2Av^810^* double mutant, mutation of *su(Hw)* in the *His2Av* null background was not enough to overcome the completely penetrant lethality associated with the inability to express the essential H2Av histone, with double mutants showing 100% lethality before the end of the pupal stage (data not shown).

Development of the larval lymph gland is influenced by *Antennapedia (Antp)*, which is expressed in the posterior signaling center (PSC), a group of cells at the posterior end of the primary lymph gland that regulates cell fate and differentiation among undifferentiated prohemocytes through paracrine signaling (Krzemień et al., 2007; Mandal et al., 2007). One possibility is that interaction of Su(Hw) and H2Av regulates the expression of *Antp* in the PSC. To address this possibility, we used publicly available ChIP-seq data to map insulator proteins and phosphorylated H2Av in the *Antp* locus. This revealed that insulator proteins Su(Hw), CP190, dCTCF, and yH2Av are enriched at the *Antp* gene, between the transcription start site and the start codon (Figure 8 C). This site does not map near the *Antp* promoter, which suggests Su(Hw) is not functioning as a transcription factor at the promoter and that *Antp* expression in the lymph gland may be regulated by a Su(Hw) insulator. We therefore wondered what the effect would be on lymph gland structure and crystal cell production when overexpressing Su(Hw) in cells that normally express *Antennapedia* during larval development. Expression of a UAS-Su(Hw)::GFP construct driven by *Antp*-GAL4 resulted in significant lethality at early developmental stages (data not shown), likely because of pleiotropic effects of the Su(Hw) overexpression, but produced larvae with severe growth defects and small melanotic tumors (Figure 8 A, B). All together, these data suggest Su(Hw) and phosphorylated H2Av both have an inhibitory effect in the developmental pathway that controls the differentiation of hemocytes in the lymph gland of *Drosophila* larva (Figure 8 D).

**Figure 8.**
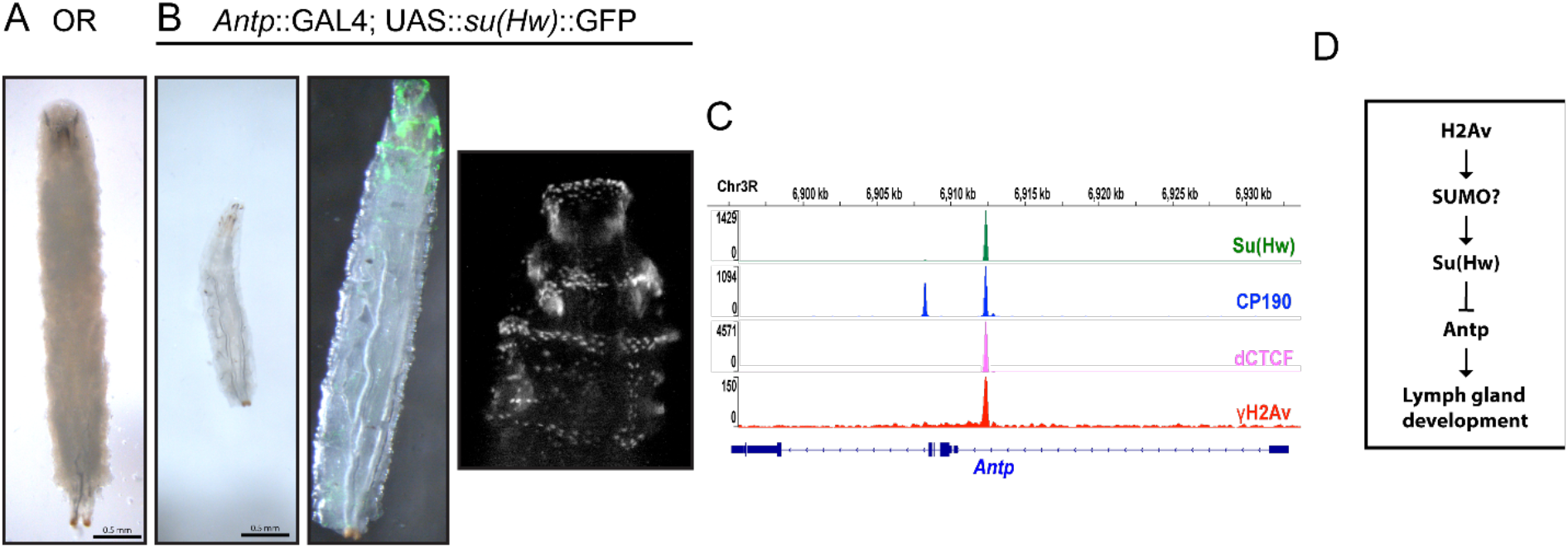
Overexpression of *su(Hw)* in *Antp*-expressing cells results in spontaneous melanization and lethality. **A.** Wild type OR larvae for size comparison. **B.** Example of *Antp*::GAL4; UAS-*su(Hw)*::GFP larvae. Left: shown at the same scale as **A**. Middle: close-up showing brightfield image overlaid with fluorescence from Su(Hw)::GFP observed at the anterior and posterior ends. Spontaneous melanization is observed on the left side of the larvae. Right: Magnified view of the anterior end of the larvae showing only Su(Hw)::GFP. The scale bars in **A.** and **B.** represent 0.5 mm. **C.** ChIP-seq data shows high enrichment of Su(Hw) insulator and other insulator proteins, including γH2Av, between the transcription start site and start codon of Antennapedia (*Antp*). **D.** Proposed model for signaling networks demonstrating the interplay between Su(Hw) and H2Av in primary lymph gland development

## Discussion

Chromatin insulators play an essential role in setting up and maintaining genome architecture during animal development by establishing long-distance interactions. These interactions help preserve the integrity of the genome against the threat of genotoxic stress and regulate the expression of genes in response to stress or developmental cues, but the mechanisms that link these activities with insulator function is not fully understood. Our previous research revealed an interaction between the Su(Hw) insulator complex and the H2Av histone variant, but the nature of the interaction and whether this was involved in maintaining genome integrity in response to DNA damage or in transcriptional regulation was unclear.

Here, we have demonstrated that the Su(Hw) insulator complex is involved in the DNA damage response. This observation is supported by our previous findings showing that null mutation of *su(Hw)* results in high levels of DNA damage during oogenesis and in the generation of gross chromosomal aberrations in dividing larval neuroblasts (Hsu et al., 2020). This mutant phenotype is common among genes that are involved in detection, marking, and resolution of double strand breaks, including the Aurora A kinase (*aurA*), ATR (*mei-41*), ATM (*tefu*), Twins (*tws*, a PP2A specificity factor), and the topoisomerase gene *Top2* (Gatti and Goldberg, 1991; Mengoli et al., 2014; Merigliano et al., 2017; Merigliano et al., 2019). This fits in with a model in which Su(Hw) also helps regulate the cellular response to DNA damage.

To test this model, larvae were subjected to UV and X-ray irradiation, which cause different types of DNA damage. Our work shows a significant enrichment increase of Su(Hw) and Mod(mdg4)67.2, an interacting partner of Su(Hw), on polytene chromatin after DNA damage. Despite this, a null mutation of *su(Hw)* significantly reduced the degree of H2Av phosphorylation in response to DNA damage by either UV or X-ray irradiation. This suggests that Su(Hw) is involved in the DNA damage response at the early step of labeling double strand breaks with H2Av phosphorylation regardless of the source of damage. Su(Hw) enrichment after UV treatment may reflect a mechanism that depends on insulator activity to respond to issues with transcription and replication that arise from pyrimidine dimer formation that are absent in the damage seen with ionizing radiation. The reliance on Su(Hw) for H2Av phosphorylation was not conserved in diploid cells of the larval central nervous system, which express little Su(Hw). These cells are still able to phosphorylate H2Av X-ray irradiation even in the absence of Su(Hw) (Figure 3), suggesting that another mechanism may regulate these processes in neurons.

Our findings on H2Av phosphorylation in polytene cells (Figure 4) also conflict with findings from meiotic cells in *Drosophila*, which show more H2Av phosphorylation in the absence of ATM/*tefu* but not ATR/*mei-41* (Joyce et al., 2011). Other results in mitotic neural cells show decreased frequency of γH2Av foci formation in the *tefu^atm-6^* mutant but not the *mei-41^29D^* mutants (Merigliano et al., 2017). The reason for these discrepancies may be due to the difference in cell types, as meiotic cells, mitotic cells, and polytene cells undergo significantly different developmental programs. The increase in Su(Hw) binding to polytene chromatin in the ATR/*mei-41^D5^* ATM/*tefu^atm-3^*double mutant would imply that more Su(Hw) accumulates in chromatin as DNA damage accrues in the absence of these two crucial DNA damage sensors. One possible mechanism is that the ATR and ATM kinases may regulate the activity of a phosphatase needed to diminish the amount of H2Av phosphorylation as DNA damage repair proceeds.

Our investigation into the relationship between H2Av and Su(Hw) further revealed significant developmental effects in larval hematopoiesis. That rescue of the melanotic tumor phenotype occurs in a Su(Hw) dose-dependent manner is not surprising, as hematopoietic phenotypes in the *His2Av^810^* mutant result from either reduction or increases in *His2Av* expression. For example, overexpression of wild type H2Av results in less cohesion between cells in the posterior signaling center (PSC, a stem cell/tissue organizing hub) and fewer differentiated plasmatocytes and crystal cells. This would imply that a precise concentration of H2Av is needed for normal development of this critical tissue. The same is likely true for Su(Hw), which regulates expression of crucial genes through stoichiometric interactions with related proteins.

The finding that single *su(Hw)^e04061^* mutants show more crystal cell clusters than in wildtype (Figure 7) implies an H2Av-independent role for Su(Hw) in crystal cell development. An H2Av-dependent function with lymph gland formation is also evident as mutation of *su(Hw)* alone is insufficient to produce the lymph gland overgrowth phenotype. Indeed, mutation of *His2Av* alone results in diminished lymph glands, a finding consistent with previous work (Grigorian et al., 2017), whereas mutation of both *su(Hw)* and *His2Av* together generates overgrown lymph glands (Figure 7 D).

An alternative interpretation is that the melanotic mass phenotype of the H2Av mutant actually depends on the phosphorylated form of H2Av, which in turn depends on Su(Hw) function, as we have previously demonstrated (Simmons et al., 2022). However, evidence is also provided in a previous study that phosphorylation of H2Av is not required for its activity in regulating hematopoietic tissue development (Grigorian et al., 2017). First, expression of a dominant negative mutant allele of *His2Av* lacking the C-terminal 14 amino acid H2AX-like motif (H2AV^CT^) (Clarkson et al., 1999) partially rescues hematopoiesis in the *His2Av^810^* null background and allows larvae to survive to adulthood (Grigorian et al., 2017). Second, overexpression of either the phosphomimetic H2AV^SE^ or the phosphonull H2AV^SA^ alleles was able to partially rescue the phenotype seen in the *His2Av^810^* null background (Grigorian et al., 2017). These results from the *His2Av^810^* background may also extend to the role that Su(Hw) is playing in this process. Su(Hw) is required for genomic stability but can also act as a transcription regulator, leading to the question of which role for Su(Hw) is involved in the hematopoietic pathway. H2Av negatively regulates the immune response in flies through modulation of another transcription factor named Relish. This is accomplished through stabilizing the levels of SUMOylation on Relish, preventing its cleavage and transcriptional activation of downstream genes (Tang et al., 2021). Whether similar regulation is involved in the cooperative activity between H2Av and Su(Hw) remains to be tested. Preliminary experiments suggest that Su(Hw) is sumoylated (data not shown), providing a possible mechanism linking the activity of H2Av to that of Su(Hw) in transcriptional regulation. Future experiments will determine if Su(Hw) is cleaved in the absence of sumoylation and the effect of sumoylation state on the transcriptional activity of Su(Hw).

One possibility to explain the influence of Su(Hw) on the H2Av-dependent development of larval lymph glands (Figure 8 D) is that Su(Hw) influences the expression of *Antp* in cells of the Posterior Signaling Center (PSC), which has downstream implications through the hedgehog signaling pathway that helps dictate cellular differentiation in the primary lobes. Particularly intriguing is the presence of an insulator in the *Antp* gene, which is enriched in both Su(Hw) and γH2Av. This insulator may function as a boundary that controls the activity of specific enhancers on the *Antp* promoter. The boundary activity may be sensitive to the expression levels of Su(Hw) and H2Av (Figure 8). Our data shows that an excess of Su(Hw) is associated with growth defects, while lack of Su(Hw) in an *His2Av* mutant background causes severe overgrowth of lymph glands (Figure 7). It is possible that overexpression of Su(Hw) in *Antp*-expressing cells causes growth defects and spontaneous melanization if Su(Hw) is necessary for regulating expression of *Antp* in cells of the PSC. Alternatively, Su(Hw) may act as a transcriptional repressor (Melnikova et al., 2019; Soshnev et al., 2013) and lack of Su(Hw) in a *His2Av* mutant background may drive aberrant expression of genes involved in stem cell maintenance in prohemocytes, which, in turn, causes overgrowth of lymph glands. Future experiments will investigate the transcriptional output of *Antp* and related genes in these mutant genotypes. An alternative hypothesis is that as the generation of reactive oxygen species (ROS) in cells of the PSC is involved in the lamellocyte maturation pathway (Sinenko et al., 2011), ROS-induced DNA damage may be an important factor in the phenotypes we described. If so, this provides another potential mechanism through which Su(Hw) may impact hematopoietic development. This remains an open question that will be addressed in future experiments.

We show here that Su(Hw) is essential for maintaining genome stability and we have characterized its relationship with the sole H2A variant in *Drosophila*. We previously demonstrated that Su(Hw) and γH2Av colocalize throughout the genome and that *His2Av* contributes to *gypsy* insulator function in chromatin and in insulator bodies (Simmons et al., 2022). We observe significantly less H2Av phosphorylation in the absence of Su(Hw); however, lack of H2Av has little effect on the recruitment or stability of insulator proteins in chromatin. From these findings we propose a model in which phosphorylation of H2Av is stabilized by an intact insulator complex and that this contributes to the function of the insulator itself. Kinase and phosphatase activity targeted to H2Av may act to regulate the activity of insulator complexes. Perhaps the most visceral manifestation of this interaction is the suppression of the melanotic tumor phenotype seen in *His2Av* mutants. These tumors are thought to result from the lack of transcriptional regulation brought about by *His2Av* mutation and does not seem related to the DNA damage response roles of H2Av (Grigorian et al., 2017). The developmental effects of mutating *su(Hw)* in the larval lymph gland in addition to *His2Av* may therefore be unrelated to the DNA damage response, although this possibility has not been ruled out. We conclude that Su(Hw) may be influencing the expression of genes in the same pathway as H2Av, and that mutation of *su(Hw)* and *His2Av* restores the proper expression program in the lymph gland, thereby preventing tumor formation. Future experiments will seek to unravel the molecular mechanism behind this interaction.

## Acknowledgements

We would like to thank Bright Amankwaa for discussions and critical review of the manuscript. Stocks obtained from the Bloomington *Drosophila* Stock Center (NIH P40OD018537) were used in this study. This work was supported partially by US Public Health Service Award from the National Institutes of Health (MH108956) with additional support from the College of Arts and Sciences and the Department of Biochemistry and Cellular and Molecular Biology at The University of Tennessee, Knoxville.

## Author contributions

Conceptualization: J.R.S., M.L.; Data curation: J.R.S.; Formal analysis: J.R.S.; Funding acquisition: M.L.; Investigation: J.R.S., J.K., M.L.; Methodology: J.R.S.; Project administration: M.L.; Resources: J.R.S., M.L.; Software: J.R.S.; Supervision: M.L.; Validation: J.R.S.; Visualization: J.R.S., M.L.; Writing – original draft: J.R.S.; Writing – review & editing: J.R.S., M.L.

---

**Supplementary Figure 1.**
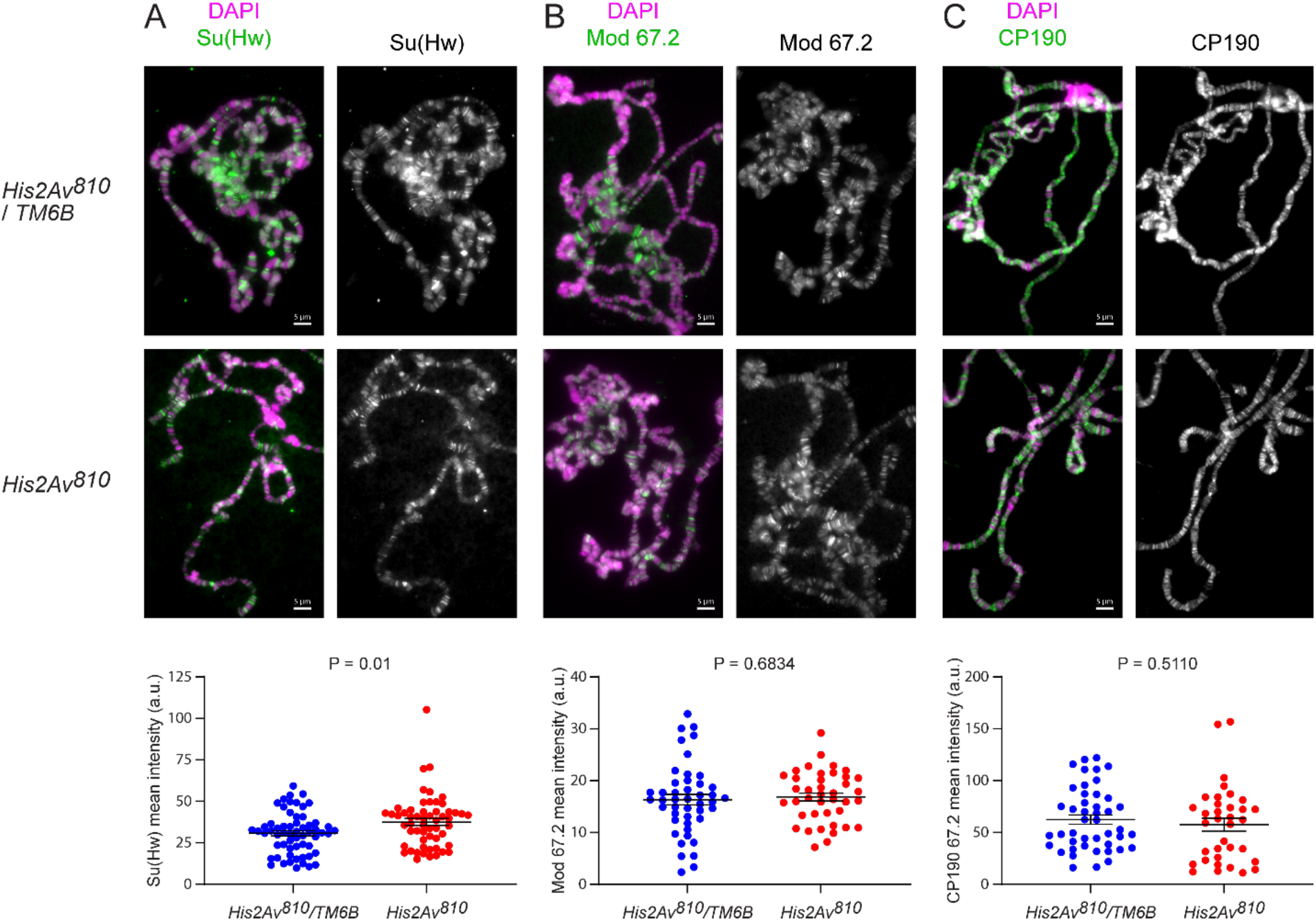
Mutation of *His2Av* has a limited effect on insulator protein binding. Immunostaining results for **A.** Su(Hw), **B.** Mod(mdg4)67.2, and **C.** CP190 antibodies on polytene chromosomes. The scale bars are 10 μm each. Beneath each figure are graphs showing fluorescent intensities of the antibody signals. Each dot represents a single polytene genome. P-values were determined using unpaired Welch’s T-tests.

**Supplementary Figure 2.**
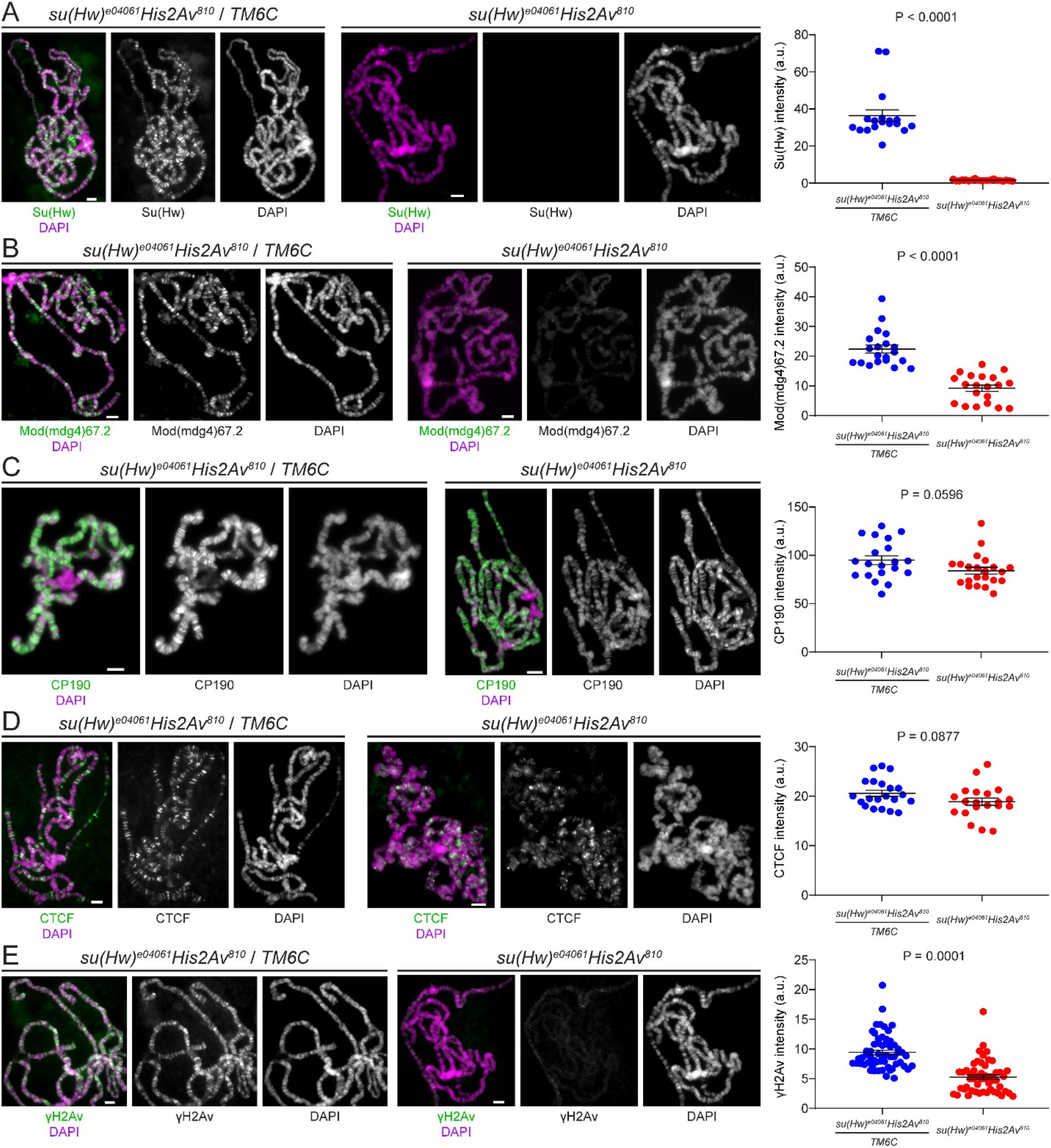
Double mutants show reduced immunostaining of Su(Hw), Mod(mdg4)67.2, and γH2Av. Immunostaining results for **A.** Su(Hw), **B.** Mod(mdg4)67.2, **C.** CP190, **D.** CTCF, and **E.** γH2Av antibodies on polytene chromosomes in *His2Av^810^* heterozygotes (left) and homozygotes (right). The scale bars are 10 μm each. To the right of each figure are graphs showing fluorescent intensities of the antibody signals. Each dot represents a single polytene genome. P-values were determined using unpaired Welch’s T-tests.

**Supplementary Figure 3.**
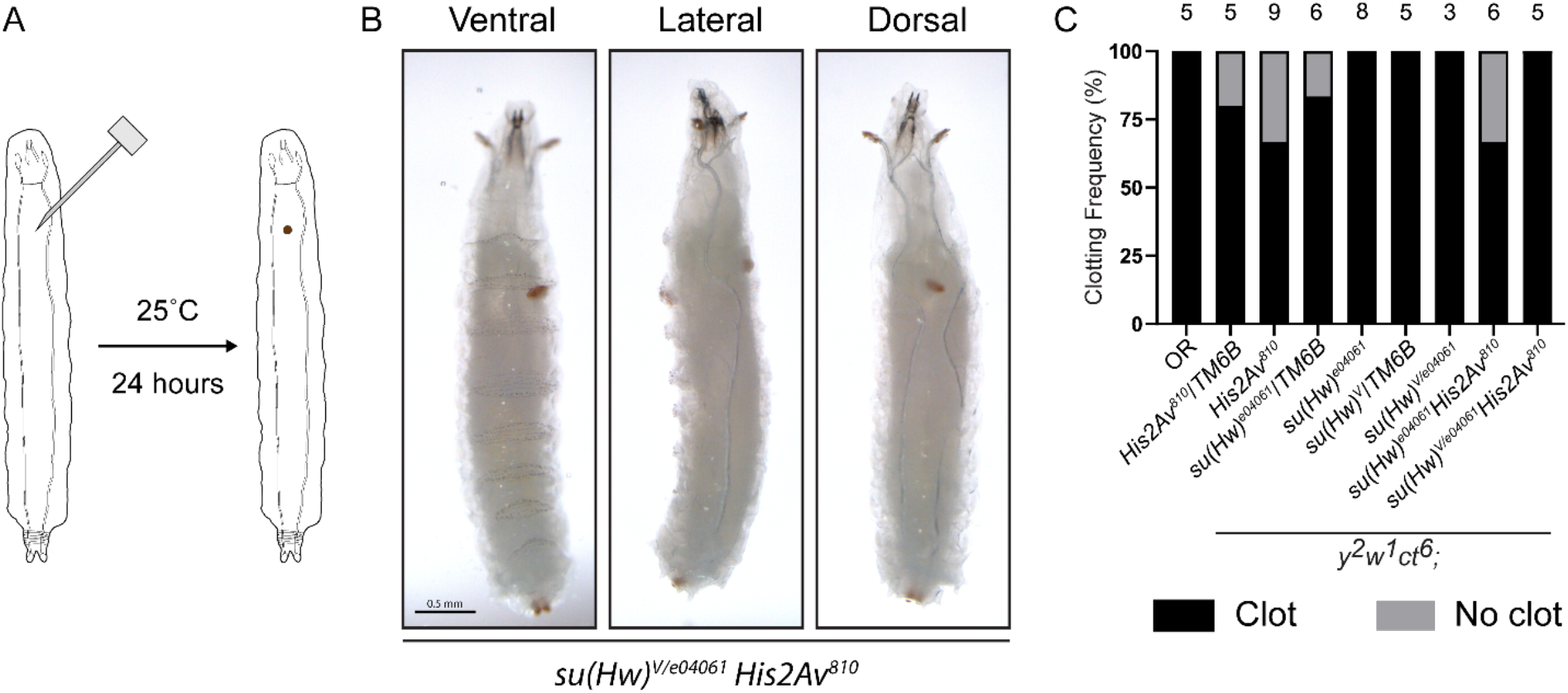
*su(Hw)^-^His2Av^810^*larvae are immunocompetent. **A.** Illustration showing the sterile wound assay, which involves stabbing larvae with a fine-tipped needle and monitoring for clot formation. **B.** Representative example of clot formation in the *y^2^w^1^ct^6^*; *su(Hw)^V/e04061^His2Av^810^* mutant genotype. Views are shown from the ventral, lateral, and dorsal aspects of the same larvae. Two clots are visible, one from the entry wound (dorsal) and one from the exit wound (ventral). **C.** Frequencies of clot formation for different genotypes.

